# HIV-1 envelope glycan loss links receptor adaptation and innate immune escape in SHIVs

**DOI:** 10.64898/2026.10.12.758606

**Authors:** Anna C. Smith, Haidyn Weight, Revansiddha H. Katte, Deepthi Kappala, Anuja Pimplapure, Vidya Mangala Prasad, Maolin Lu, Julie Overbaugh, Amit Sharma

## Abstract

Simian-human immunodeficiency viruses (SHIVs) are widely used to model HIV-1 infection in macaques, yet most SHIVs encoding circulating HIV-1 envelope glycoproteins (Env) replicate poorly because of inefficient use of macaque CD4 and sensitivity to host antiviral responses. During adaptation to macaques, Env is therefore subject to multiple selective pressures to improve receptor utilization while maintaining function in the face of host immune restriction. Type I interferon (IFN) is a major component of the host antiviral response during acute infection, and IFN resistance increases during SHIV adaptation to macaques through serial passage. However, the viral determinants of this resistance have remained unclear. Here, we show that serial passage of a lineage of SHIVs in macaques selected Env variants with markedly increased resistance to IFN-α and enhanced replication in macaque CD4^+^ T lymphocytes. IFN-α resistance mapped principally to loss of two highly conserved N-linked glycans at N197 and N276 near the CD4-binding site. Glycan loss altered the conformational landscape of virus-associated Env, enhanced macaque CD4-dependent entry and membrane fusion without increasing human CD4-dependent entry, and reduced incorporation of the IFN-inducible restriction factors IFITM3 and IFITM3A into progeny virions. Depletion of IFITM3/3A rendered the parental SHIV highly resistant to IFN-α and abolished differential IFN-α sensitivity among parental, adapted, and glycan-loss viruses. These findings reveal that loss of receptor-proximal Env glycans can simultaneously improve utilization of a heterologous CD4 receptor and promote escape from IFN-mediated restriction, linking receptor adaptation and innate immune evasion during lentiviral adaptation to a new host.

**Significance:** Simian-human immunodeficiency viruses (SHIVs) are essential for modeling HIV-1 infection in macaques, but their adaptation to macaques requires HIV-1 envelope glycoprotein (Env) to overcome species-specific barriers to replication. We show that loss of two conserved Env glycans selected during macaque passage simultaneously improves macaque CD4 usage and promotes escape from IFITM-mediated restriction. These findings reveal that receptor incompatibility and innate immune restriction can impose convergent selective pressures on Env, allowing receptor adaptation and innate immune escape to arise through the same Env changes rather than independent evolutionary solutions. More broadly, they identify the HIV-1 glycan shield as a functional interface linking receptor usage, membrane fusion, and resistance to host antiviral defenses during cross-species adaptation.

## Introduction

Simian-human immunodeficiency viruses (SHIVs), which express HIV-1 envelope glycoproteins (Env) in an SIV backbone, are widely used in macaques to model HIV-1 transmission, pathogenesis, and antibody-mediated protection (1). Because Env is the sole target of HIV-1 neutralizing antibodies, SHIVs are therefore particularly valuable for evaluating vaccine- and antibody-mediated protection against infection. Ideally, SHIV challenge viruses should encode Envs representative of circulating or transmitted/founder (T/F) HIV-1 variants (2). However, most SHIVs encoding such Envs replicate poorly and require adaptation through serial passage in macaques to achieve robust replication and establish persistent infection (3–9). A major determinant of this species barrier is the inefficient use of macaque CD4 for viral entry by most circulating and T/F HIV-1 Envs (10). Accordingly, adaptation of SHIV Env to macaque CD4, both in vitro and in vivo, can select substitutions that enhance CD4 engagement, alter Env conformation, or both (11–13). Such adaptations can increase SHIV replication but may also alter Env antigenicity, potentially limiting how faithfully adapted SHIVs recapitulate the biological properties of circulating and T/F Envs (12).

Receptor incompatibility is not the only host barrier encountered during SHIV adaptation. Because Env must retain its essential entry function while adapting to a non-native CD4 receptor, it is subject to strong selective pressure during replication in macaques. Env is also shaped by host humoral immune pressure, further constraining the adaptive pathways available to the virus (14–16). In addition to these receptor and humoral immune pressures, the type I interferon (IFN) response generates a potent antiviral state through induction of numerous interferon-stimulated genes, including host restriction factors that inhibit multiple stages of lentiviral replication (17–19). Acute HIV-1 and SIV infections are accompanied by robust type I IFN responses during the early stages of infection (19–22). Consistent with this, resistance to type I IFNs has been associated with increased viral fitness during early HIV-1 infection (23, 24).

We previously found that unadapted SHIVs encoding circulating HIV-1 Envs are more sensitive to IFN-α than macaque-adapted SHIVs and that serial passage in macaques is accompanied by increasing IFN-α resistance (25). These observations suggest that the IFN response may impose an additional selective pressure during adaptation of HIV-1 Env to the macaque host, but the viral determinants underlying the acquisition of IFN-α resistance during serial passage have remained undefined.

Interferon-induced transmembrane (IFITM) proteins represent a potential mechanistic link between Env function and the IFN-induced antiviral state. IFITMs are membrane-associated restriction factors that inhibit infection of diverse enveloped viruses, in part by interfering with viral-host membrane fusion (26, 27). IFITMs can also be incorporated into HIV-1 particles, where they impair particle infectivity and fusion (28, 29). Susceptibility to IFITM restriction varies among viral glycoproteins, indicating that glycoprotein properties can influence IFITM-mediated inhibition (30–33). The macaque IFITM repertoire has been expanded through lineage-specific gene duplication and retrotransposition and includes canonical IFITM1 and IFITM3, the duplicated paralog IFITM3A, and two IFITM3-derived retrocopies, IFITM3-R1 and IFITM3-R2 (34). We previously showed that incorporation of macaque IFITMs into SHIV particles is Env dependent and that macaque IFITMs preferentially restrict SHIVs encoding unadapted HIV-1 Envs (34). In contrast, macaque-adapted SHIVs are comparatively resistant to both IFN- and IFITM-mediated restriction (25, 34). Together, these observations raise the possibility that Env properties governing adaptation to macaque CD4 may also influence susceptibility to IFN-induced restriction. Whether receptor adaptation and escape from IFN-mediated restriction represent independent adaptive processes or can arise through common changes in Env has remained unclear.

Here, we define the viral determinants underlying the acquisition of IFN-α resistance during serial passage of a lineage of SHIVs in macaques. We show that passage-associated IFN-α resistance maps to Env and is conferred principally by loss of two highly conserved N-linked glycans at N197 and N276 near the CD4-binding site. Loss of these glycans alters the Env conformational landscape, enhances macaque CD4-dependent entry and membrane fusion, and reduces incorporation of macaque IFITM3 and IFITM3A into progeny virions. Depletion of endogenous IFITM3/3A abolishes the differential IFN-α sensitivity of parental and adapted viruses, establishing these restriction factors as major determinants of the Env-dependent phenotype. Thus, passage-associated glycan loss mechanistically links adaptation to a heterologous CD4 receptor with escape from IFN-mediated restriction, revealing how a common set of Env adaptations can overcome two distinct barriers to lentiviral replication in a new host.

## Results

### Serial macaque passage of SHIV selects Env-mediated resistance to IFN-α

Adaptation of SHIVs to macaques commonly involves serial passage to increase viral replication and pathogenicity. Rapid serial passage is performed within the first two weeks of infection, a time during which macaques mount a robust type I IFN response (19–21). We previously showed that serial passage increases SHIV resistance to IFN-α (25). However, the viral determinants of IFN-α resistance in serially passaged SHIVs have not been defined. To identify the viral adaptations associated with enhanced replication and IFN-α resistance, we examined a lineage of SHIVs derived by sequential macaque passage of SHIV-109mc (35). SHIV-109mc encodes the HIV-1 ZM109F.PB4 Env (36), an early-stage subtype C Env derived directly from patient PBMCs without culture adaptation and included in the subtype C reference Env panel used for assessment of vaccine-elicited neutralizing antibodies (37). Virus populations recovered from the passage 3 animal at 24 weeks postinfection (SHIV-P3) and at necropsy at 34 weeks postinfection (SHIV-P3N), as well as from the passage 4 animal at 22 weeks postinfection (SHIV-P4), were compared with parental SHIV-109mc (35) (Fig. 1A).

**Figure 1.**
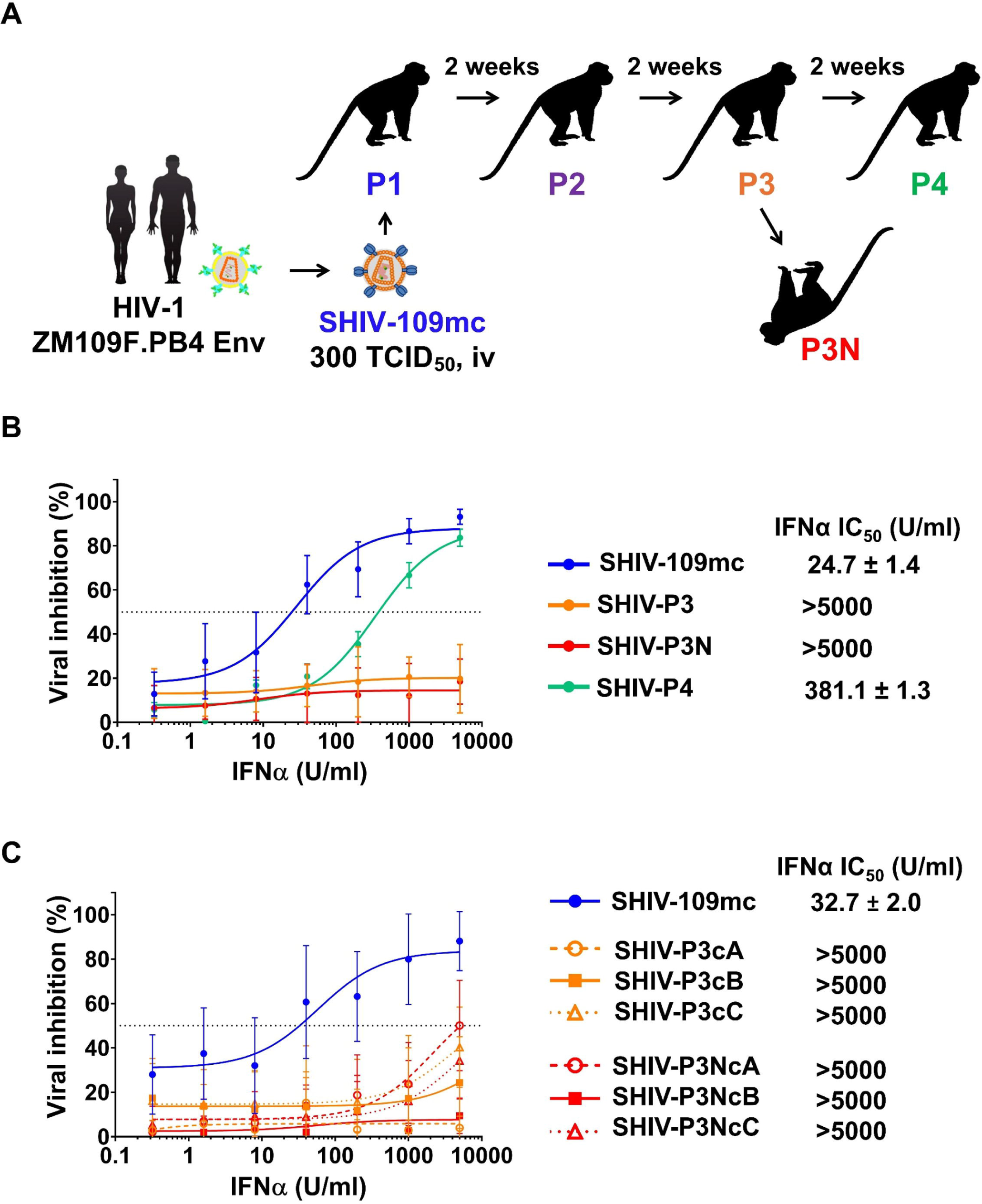
**Serial macaque passage of SHIV selects Env-mediated resistance to IFN-**α**. (A)** Experimental scheme for *in vivo* adaptation of SHIV-109mc, which encodes the subtype C HIV-1 ZM109F.PB4 Env. Parental SHIV-109mc was inoculated intravenously into rhesus macaques and transferred at 2-week intervals through four sequential passages. Virus populations recovered from the passage 3 animal at 24 weeks postinfection (SHIV-P3) and at necropsy at 34 weeks postinfection (SHIV-P3N), as well as from the passage 4 animal at 22 weeks postinfection (SHIV-P4), were analyzed. **(B)** IFN-α dose-response profiles of parental SHIV-109mc and the indicated macaque-passaged virus populations in immortalized macaque CD4^+^ T lymphocytes. Cells were infected at an MOI of 0.02 and cultured with increasing concentrations of IFN-α. Viral infectivity at 7 days postinfection was quantified using TZM-bl indicator cells and expressed as percent inhibition relative to matched untreated cultures. **(C)** IFN-α dose-response profiles of infectious molecular clones encoding Env sequences recovered from the P3 or P3N virus populations. Curves were fitted by nonlinear regression; the dotted gray line indicates 50% inhibition, and calculated IFN-α IC_50_ values are shown. Data are means from three independent experiments performed in duplicate; error bars indicate standard deviations.

We first compared the IFN-α sensitivity of the parental and passaged virus populations in immortalized macaque CD4^+^ T lymphocytes. SHIV-109mc was potently inhibited by IFN-α, with a mean IC_50_ of ∼25 U/mL. By contrast, SHIV-P4 exhibited approximately 15-fold greater resistance (mean IC_50_ ∼381 U/ml), whereas SHIV-P3 and SHIV-P3N remained resistant even at the highest IFN-α concentration tested (IC_50_ >5,000 U/ml) (Fig. 1B). Notably, the observed IFN-mediated restriction in the immortalized macaque CD4^+^ T lymphocytes was comparable to that observed previously in primary rhesus macaque lymphocytes (25). Thus, consistent with our previous findings, sequential passage in macaques selected viruses with markedly reduced sensitivity to IFN-α.

Because HIV-1 Env can influence susceptibility to IFN-induced restriction (25, 34), we next asked whether the resistance selected during passage mapped to Env. Phylogenetic and sequence analyses of full-length Env clones recovered from the P3 and P3N virus populations showed related but genetically diverse lineages, with >95% overall sequence identity among the isolates (Fig. S1). Compared with SHIV-109mc Env, passage-derived Envs accumulated substitutions predominantly within the C2 and C3 constant regions and the V3 and V4 variable loops of gp120 (Fig. S2).

We next generated the parental SHIV-109mc infectious molecular clone (IMC) and six lineage-related IMCs by introducing three independent P3 Envs (P3cA, P3cB, and P3cC) and three independent P3N Envs (P3NcA, P3NcB, and P3NcC) into a common SIVmac239-derived SHIV1157ipd3N4 backbone (8), generating viruses that differed only in Env sequence. All six IMCs encoding passage-derived Envs were highly resistant to IFN-α, with IC_50_ values >5,000 U/ml, whereas SHIV-109mc remained strongly IFN-α sensitive (mean IC_50_ ∼33 U/ml; Fig. 1C). These findings demonstrate that passage-associated changes in Env are sufficient to confer IFN-α resistance.

The passage-derived Env IMCs also replicated more efficiently than SHIV-109mc in macaque CD4^+^ T lymphocytes (Fig. S3A). Across the virus panel, replication capacity strongly correlated with IFN-α resistance (Spearman *r*_s_ = 0.9870, *P* < 0.0001; Fig. S3B), linking the Env-dependent IFN-α-resistant phenotype with increased viral fitness in macaque cells. Collectively, these data indicate that serial passage in macaques selected multiple independently evolved Env variants that conferred enhanced replication and robust resistance to IFN-α.

### IFN-α resistance maps to gp120 and is conferred by loss of the N197 and N276 glycans

We next sought to identify the Env determinants responsible for IFN-α resistance. Because SHIV-P3cB exhibited the strongly IFN-α-resistant phenotype characteristic of the passage-derived Envs and its Env was genetically closest to 109mc Env, we generated a series of Env chimeras between SHIV-109mc and SHIV-P3cB (Fig. 2A). These chimeras introduced the entire gp120 surface subunit, the gp41 transmembrane subunit, or the gp120 C2-V4 region containing most passage-associated substitutions. Transfer of P3cB gp120 into SHIV-109mc conferred complete resistance to IFN-α (IC_50_ >5,000 U/ml), whereas replacement of gp41 did not (mean IC_50_ 0.6 U/ml), localizing the dominant determinant to gp120 (Fig. 2B). A chimera containing the P3cB C2-V4 region was similarly resistant (IC_50_ >5,000 U/ml), further narrowing the determinant to this region of gp120.

**Figure 2.**
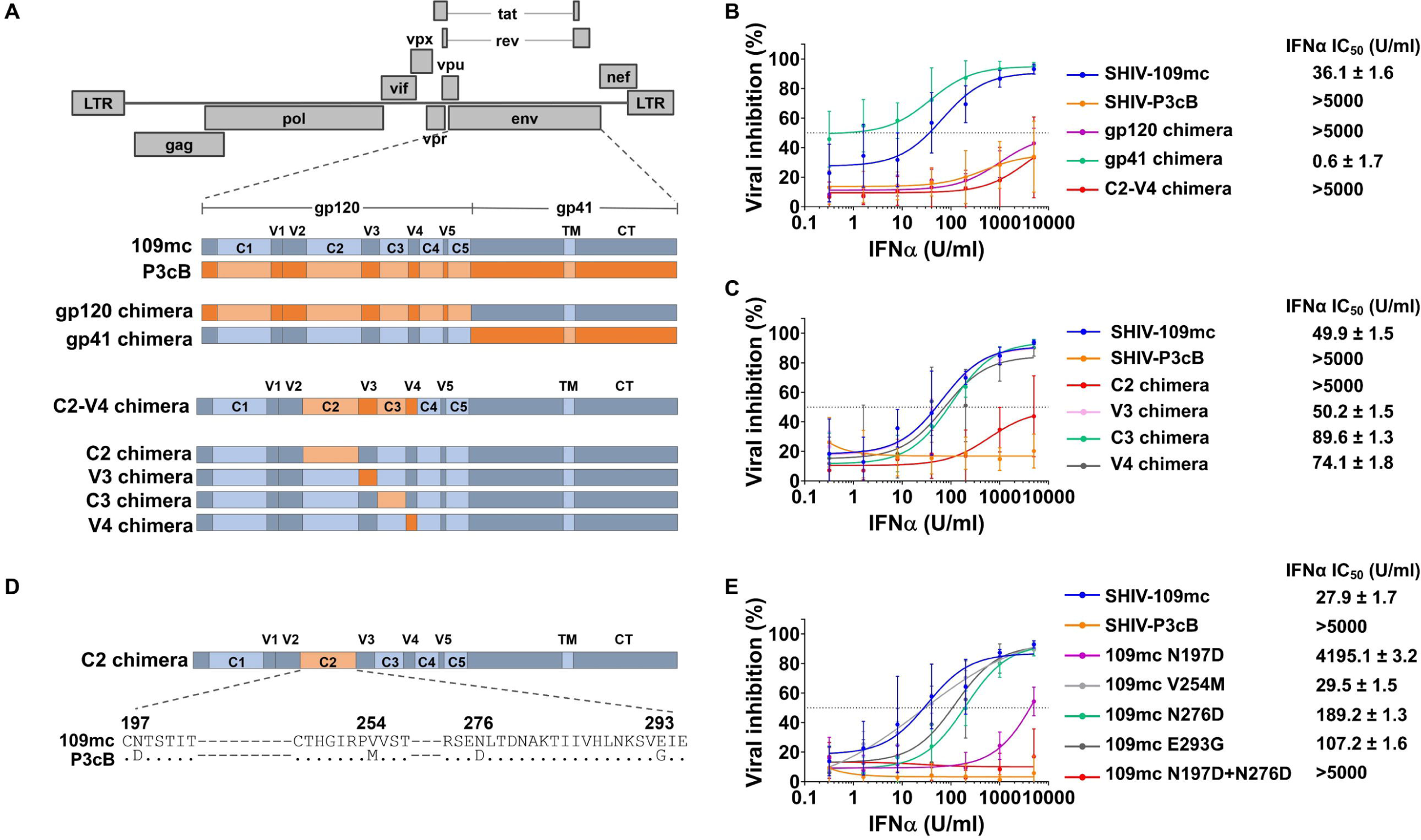
**Loss of N-linked glycans at N197 and N276 confers SHIV resistance to IFN-**α**. (A)** Schematics of the SHIV genome and Env chimeras generated between IFN-α-sensitive SHIV-109mc (blue) and IFN-α-resistant SHIV-P3cB (orange). Chimeras exchanged gp120, gp41, the C2-V4 region, or the individual C2, V3, C3, and V4 subregions. **(B)** IFN-α dose-response profiles of SHIV-109mc, SHIV-P3cB, and SHIVs encoding the gp120, gp41, or C2-V4 chimeric Envs. **(C)** IFN-α dose-response profiles of SHIVs encoding the indicated gp120 subregion chimeras. **(D)** Alignment of SHIV-109mc and SHIV-P3cB Env sequences highlighting the N197D, V254M, N276D, and E293G substitutions. Dots indicate amino acid identity. **(E)** IFN-α dose-response profiles of SHIV-109mc encoding the indicated individual substitutions or the combined N197D+N276D substitutions. Curves were fitted by nonlinear regression; the dotted gray line indicates 50% inhibition, and calculated IFN-α IC_50_ values are shown. Data are means from three independent experiments performed in duplicate; error bars indicate standard deviations.

To further map the determinant within the C2-V4 region of gp120, individual C2, V3, C3, and V4 subregions were transferred from P3cB into SHIV-109mc (Fig. 2A). Among these chimeras, transfer of the P3cB C2 region was sufficient to reproduce the IFN-α resistance of SHIV-P3cB, with no measurable inhibition at the highest IFN-α concentration tested. In contrast, the V3, C3, and V4 chimeras remained IFN-α sensitive, with IC_50_ values of ∼50 to 90 U/ml (Fig. 2C). Compared with 109mc Env, the C2 region of IFN-α-resistant P3cB Env contained four amino acid substitutions: N197D, V254M, N276D, and E293G (Fig. 2D). Notably, N197D and N276D disrupt conserved potential N-linked glycosylation sites positioned near the CD4-binding site of gp120 (38, 39).

We therefore introduced each substitution individually into SHIV-109mc and assessed its effect on IFN-α sensitivity. V254M had little effect, whereas E293G produced a ∼4-fold increase in resistance. In contrast, N197D and N276D increased IFN-α resistance by approximately 150-fold and 7-fold, respectively (Fig. 2E). Because N197D reproduced most, but not all, of the IFN-α-resistant phenotype, we combined N197D and N276D to determine whether the substitutions had complementary effects. The N197D+N276D double mutant fully phenocopied SHIV-P3cB, with no measurable inhibition at the highest IFN-α concentration tested (Fig. 2E). Although a minor contribution of E293G cannot be excluded, the ability of N197D+N276D to fully reproduce the SHIV-P3cB phenotype identifies loss of these two glycans as the principal determinant of IFN-α resistance.

Replication analyses of the chimeric and mutant SHIVs in macaque CD4^+^ T lymphocytes showed a concordant pattern. Replication capacity closely tracked with IFN-α resistance across the gp120/gp41/C2-V4 chimera panel and the SHIV-109mc point-mutant panel (Fig. S4). The association was particularly strong for the larger-region chimeras (*r*_s_ = 0.9795, *P* = 0.0013) and point mutants (*r*_s_ = 0.9660, *P* < 0.0001), although it did not reach significance when the individual C2-V4 subregion chimeras were analyzed separately (*r*_s_ = 0.6154, *P* = 0.0647). Together, these results identify loss of the N197 and N276 glycans as major Env adaptations that enhance SHIV replication and confer resistance to IFN-α.

### Loss of N197 and N276 glycans alters Env conformational landscape and enhances macaque CD4-dependent entry and membrane fusion

The selection of N197 and N276 glycan loss during passage raised the question of how these Env changes enhance SHIV replication and IFN-α resistance in macaque cells. To examine the structural context of these residues, we generated a homology model of SHIV-109mc Env using structures of the ZM109F.PB4 gp120 core (40) and BG505 Env trimer (41) (Fig. 3A). Structural superposition showed that the positions of N197 and N276 in 109mc Env closely aligned with the corresponding glycosylated residues in BG505 Env (Fig. 3B). Modeling of gp120 in relation to human and macaque CD4 further positioned both glycans proximal to the CD4-binding site (Fig. 3C). The N197 glycan was predicted to extend into the path of CD4 engagement, whereas the N276 glycan was positioned nearby without an apparent direct steric clash. Together with published structures of deglycosylated subtype A and C Env trimers (42, 43), these observations suggest that loss of the N197 and N276 glycans could increase accessibility of the CD4-binding site.

**Figure 3.**
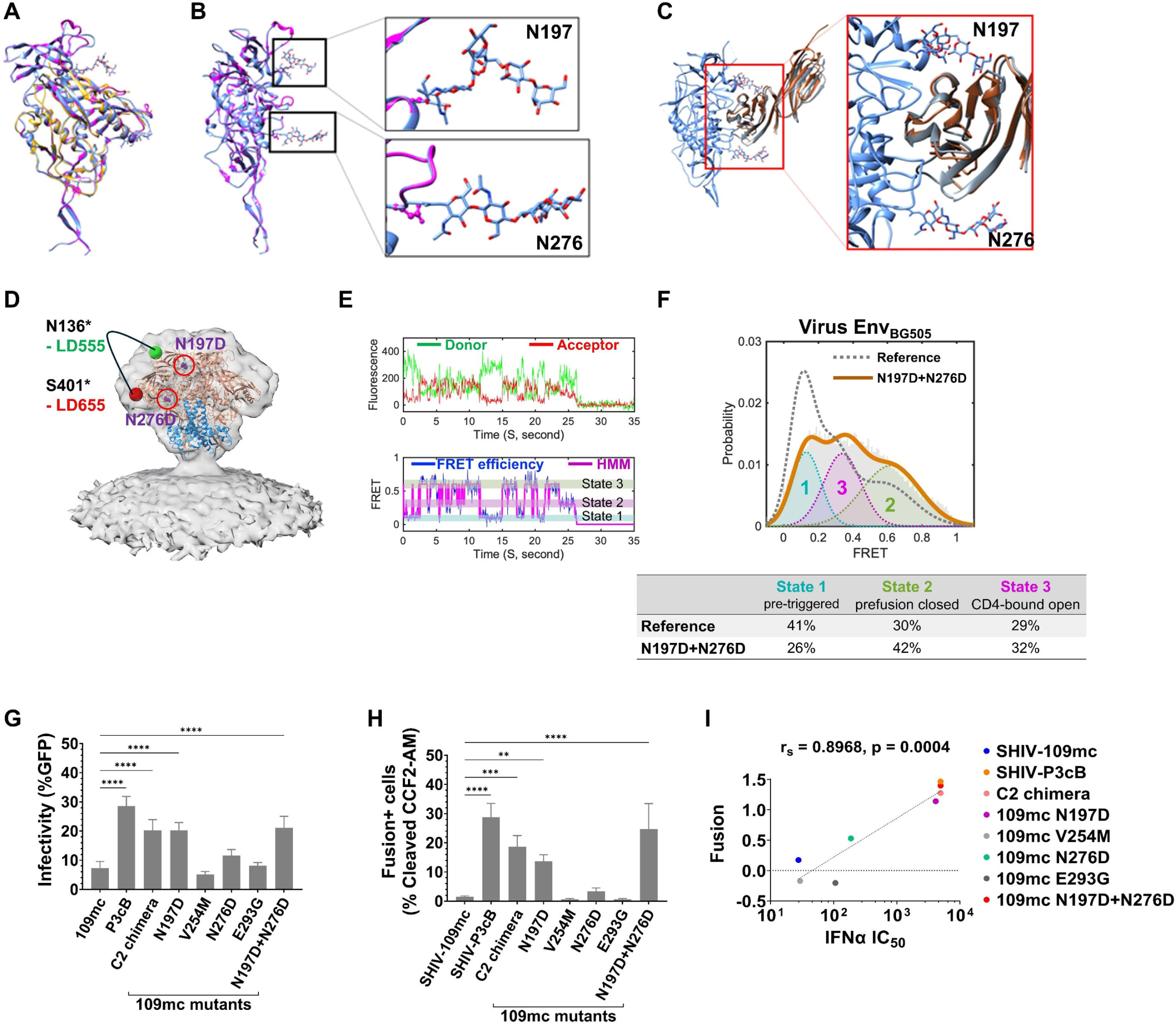
Loss of N197 and N276 glycans alters Env conformational landscape and enhances macaque CD4-dependent entry and membrane fusion. (A) Structural superposition of gp120 subunits from BG505 SOSIP.664 (blue; PDB 5FYL), ZM109F.PB4 gp120 core (yellow; PDB 3TIH), and a homology model of SHIV-109mc Env (magenta). (B) Rotated view of the superimposed BG505 SOSIP.664 and SHIV-109mc Env models showing the positions of the N197- and N276-linked glycans, with enlarged views of each glycan site. (C) Ribbon representation showing human CD4 (brown; PDB 1GC1) and macaque CD4 (gray; PDB 6TYB) in relation to gp120. Boxed region indicates the gp120-CD4 interaction interface; the enlarged view shows the positions of the N197 and N276 glycans relative to the CD4-binding interface. (D) Schematic of the single-molecule Förster resonance energy transfer (smFRET) labeling strategy used to monitor conformational dynamics of virus-associated HIV-1 BG505 Env bearing the N197D and N276D substitutions. Env trimer structure (PDB 4ZMJ) was fitted into the density map of membrane-associated Env (EMD 21412), with sites used for attachment of donor (LD555) and acceptor (LD655) fluorophores indicated within a single Env protomer. (E) Representative fluorescence intensity trajectories for donor (green) and acceptor (red) fluorophores and the corresponding FRET efficiency trace (blue) over 35 seconds. Hidden Markov model (HMM) idealization of the FRET trajectory (magenta) is shown, with the three conformational states indicated. (F) FRET efficiency distribution compiled from individual dynamic trajectories of BG505 Env bearing the N197D and N276D substitutions (*n* = 138), overlaid with the corresponding wild-type BG505 Env reference distribution (dashed gray). Histograms were fitted to a constrained three-state Gaussian model corresponding to the pre-triggered (dashed cyan), prefusion-closed (dashed red), and CD4-bound open (dashed green) conformational states. Estimated state occupancies are indicated. **(G)** Entry mediated by the indicated Envs in HEK293T cells expressing macaque CD4 and CCR5. GFP-reporter viruses pseudotyped with each Env were used to infect target cells, and entry was quantified as the percentage of GFP-positive cells by flow cytometry. Data are means from four independent experiments; error bars indicate standard deviations. Statistical significance was determined by ordinary one-way ANOVA followed by Dunnett’s multiple-comparisons test relative to 109mc Env. **(H)** Membrane fusion mediated by the indicated SHIVs. BlaM-Vpr-containing SHIV particles were normalized by reverse transcriptase activity and used to infect immortalized macaque CD4^+^ T lymphocytes. Fusion was quantified as the percentage of cells containing cleaved CCF2-AM substrate. Data are means from three independent experiments; error bars indicate standard deviations. Statistical significance was determined by ordinary one-way ANOVA followed by Dunnett’s multiple-comparisons test relative to SHIV-109mc. **(I)** Relationship between SHIV membrane fusion and IFN-α resistance. Mean fusion values were plotted against the corresponding IFN-α IC_50_ values, and the association was evaluated by Spearman rank correlation (*r*_s_ = 0.8968, *P* = 0.0004). \*\**P* < 0.01; \*\*\**P* < 0.001; \*\*\*\**P* < 0.0001.

To determine whether loss of the N197 and N276 glycans altered the conformational landscape of Env, we performed single-molecule Förster resonance energy transfer (smFRET) analysis of virus-associated Env bearing the N197D and N276D substitutions using the bioorthogonal labeling approach (44) (Fig. 3 D–F). Individual fluorescence trajectories exhibited transitions among three previously defined conformational states corresponding to a pre-triggered state (State 1), a prefusion-closed state (State 2), and a CD4-bound open state (State 3) (45, 46) (Fig. 3E). Compared with the wild-type Env reference, N197D+N276D Env displayed a redistribution of conformational state occupancies: occupancy of the pre-triggered State 1 decreased from 41% to 26%, whereas occupancy of the prefusion-closed State 2 increased from 30% to 42%. Occupancy of the CD4-bound open State 3 changed comparatively little, from 29% to 32% (Fig. 3F and Table S1). Thus, removal of the N197 and N276 glycans shifts the Env conformational ensemble away from the pre-triggered state and toward the prefusion-closed state, providing direct evidence that glycan loss alters Env conformational landscape. Together with the structural modeling, these findings indicate that loss of the N197 and N276 glycans alters both the local structural context of the CD4-binding site and the conformational landscape of Env.

We next examined the effect of glycan loss on viral entry using GFP-reporter viruses pseudotyped with the indicated Envs and HEK293T target cells expressing macaque CD4 and CCR5. Compared with parental 109mc Env, P3cB Env mediated approximately 4-fold greater entry. Introduction of the P3cB C2 region, N197D alone, or N197D+N276D into 109mc Env enhanced macaque CD4-dependent entry by approximately 3-fold, whereas N276D, V254M, and E293G did not significantly increase entry (Fig. 3G). In contrast, no significant differences were detected when the same Envs were tested in cells expressing human CD4 and CCR5 (Fig. S5), indicating that the entry advantage associated with N197 glycan loss was preferentially observed in the context of macaque CD4.

We next asked whether enhanced macaque CD4-dependent entry translated into increased membrane fusion by replication-competent SHIV. Using BlaM-Vpr-containing particles normalized for reverse transcriptase activity, we quantified fusion with macaque CD4^+^ T lymphocytes. Compared with parental SHIV-109mc, SHIV-P3cB exhibited approximately 19-fold greater membrane fusion, whereas N197D increased fusion by approximately 9-fold (Fig. 3H and Fig. S6). N276D produced a more modest ∼2-fold, nonsignificant increase, whereas V254M and E293G did not significantly alter fusion. The N197D+N276D double mutant displayed an approximately 17-fold increase in fusion, approaching the phenotype of SHIV-P3cB and consistent with the complementary effects of these substitutions on IFN-α resistance. Importantly, membrane-fusion efficiency strongly correlated with IFN-α resistance across the virus panel (Spearman *r*_s_ = 0.8968, *P* = 0.0004; Fig. 3I). Together, these findings link loss of the N197 and N276 glycans to altered Env conformational landscape, enhanced macaque CD4-dependent entry, and increased membrane fusion, thereby connecting viral entry and fusion with IFN-α resistance.

### Env glycan loss reduces virion incorporation of IFITM3/3A

We next investigated how glycan-dependent changes in Env function might contribute to resistance against the IFN-induced antiviral state. We previously showed that macaque interferon-induced transmembrane (IFITM) proteins can restrict SHIV in an Env-dependent manner and that incorporation of endogenous macaque IFITM3 and its duplicated paralog IFITM3A into SHIV particles is also determined by Env (34). We therefore asked whether serial macaque passage altered the association of IFITM3/3A with SHIV particles. Endogenous IFITM3/3A was readily detected in SHIV-109mc particles produced from macaque CD4^+^ T lymphocytes but was reduced in particles from the passaged SHIV-P3 and SHIV-P3N populations (Fig. 4A), suggesting that passage-associated Env adaptation influences the extent of IFITM3/3A incorporation into progeny virions.

**Figure 4.**
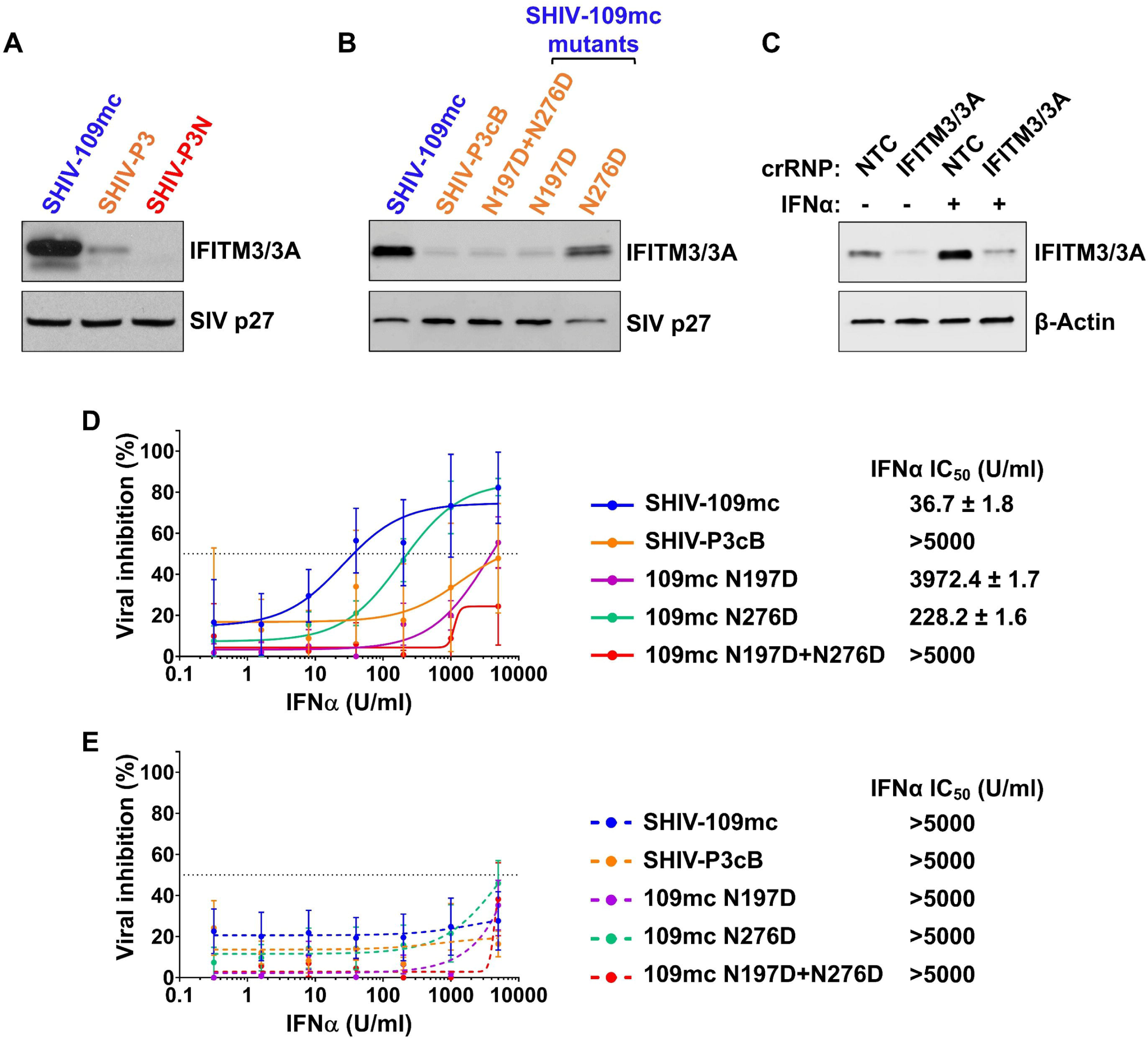
Env glycan loss reduces virion IFITM3/3A incorporation and confers resistance to IFITM-mediated restriction. **(A)** Incorporation of endogenous macaque IFITM3/3A into parental and macaque-passaged SHIV particles. Cell-free SHIV-109mc, SHIV-P3, and SHIV-P3N virions recovered from infected macaque CD4^+^ T lymphocytes were normalized by reverse transcriptase activity and analyzed by immunoblotting for macaque IFITM3/3A and SIV p27. **(B)** Incorporation of endogenous IFITM3/3A into SHIV-109mc, SHIV-P3cB, and the indicated SHIV-109mc glycan mutants. Virions were normalized and analyzed as in (A). **(C)** CRISPR-Cas9 ribonucleoprotein-mediated depletion of IFITM3/3A in immortalized macaque CD4^+^ T lymphocytes. Cells were electroporated with a nontargeting control crRNP (NTC) or an IFITM3/3A-targeting crRNP and cultured in the absence or presence of IFN-α. Cell lysates were analyzed by immunoblotting for IFITM3/3A and β-actin. **(D)** IFN-α dose-response profiles of SHIV-109mc, SHIV-P3cB, and the indicated SHIV-109mc glycan mutants in NTC-treated macaque CD4^+^ T lymphocytes. **(E)** IFN-α dose-response profiles of the same viruses in IFITM3/3A-depleted macaque CD4^+^ T lymphocytes. Curves were fitted by nonlinear regression; the dotted gray line indicates 50% inhibition, and calculated IFN-α IC_50_ values are shown. Data are means from three independent experiments performed in duplicate; error bars indicate standard deviations.

To determine whether Env was sufficient to account for the differential incorporation of IFITM3/3A into SHIV particles, we compared SHIV-109mc with the Env-reconstructed SHIV-P3cB IMC and the individual glycan-loss mutants. SHIV-P3cB particles contained less endogenous IFITM3/3A than SHIV-109mc particles despite otherwise sharing a common proviral background (Fig. 4B), indicating that differences in Env were sufficient to alter virion-associated IFITM3/3A. Introduction of N197D markedly reduced virion-associated IFITM3/3A, both alone and in combination with N276D, whereas N276D alone had a more limited effect on IFITM3/3A incorporation (Fig. 4B). This hierarchy paralleled the relative effects of these substitutions on IFN-α sensitivity, replication, macaque CD4-dependent entry, and membrane fusion.

Ectopic-expression experiments with individual macaque IFITM homologs further supported Env-dependent differences in IFITM incorporation. Although multiple macaque IFITM proteins were expressed in producer cells, their incorporation into SHIV particles differed according to both IFITM identity and Env. The most pronounced differences between SHIV-109mc and SHIV-P3cB were observed for IFITM3 and IFITM3A, which were more abundant in SHIV-109mc particles than in SHIV-P3cB particles (Fig. S7). Together, these observations indicate that Env glycan composition influences the extent of IFITM3/3A incorporation into SHIV particles and link passage-associated Env adaptation to reduced virion association with these IFN-induced restriction factors.

### IFITM3/3A depletion eliminates Env-dependent differences in IFN-α sensitivity

Finally, we asked whether IFITM3/3A accounted for the Env glycan-dependent differences in IFN-α sensitivity between parental, adapted, and glycan-loss SHIVs. Endogenous IFITM3/3A was depleted from immortalized macaque CD4^+^ T lymphocytes using CRISPR-Cas9 ribonucleoprotein complexes. Immunoblotting confirmed efficient depletion of IFITM3/3A relative to cells receiving a nontargeting control, under both basal conditions and following IFN-α treatment (Fig. 4C).

We then compared IFN-α sensitivity in control and IFITM3/3A-depleted cells. In nontargeting control cells, parental SHIV-109mc remained highly sensitive to IFN-α (mean IC_50_ ∼36 U/ml), whereas SHIV-P3cB and the SHIV-109mc N197D, N276D, and N197D+N276D mutants were substantially more resistant, with IC_50_ values comparable to those previously measured in untreated cells (Fig. 2E and Fig. 4D). In contrast, depletion of IFITM3/3A rendered parental SHIV-109mc highly resistant to IFN-α, increasing its IC_50_ from ∼36 U/ml in control cells to >5,000 U/ml (Fig. 4E). SHIV-P3cB and each of the glycan-loss mutants were similarly resistant in IFITM3/3A-depleted cells, with IC_50_ values >5,000 U/ml (Fig. 4E). Thus, depletion of IFITM3/3A abolished the differences in IFN-α sensitivity between parental SHIV-109mc and the adapted and glycan-loss viruses. These results genetically link IFITM3/3A to the Env-dependent IFN-α phenotype and demonstrate that the differential IFN-α sensitivity associated with the N197 and N276 glycans requires the IFITM3/3A restriction pathway.

## Discussion

Receptor incompatibility and innate immune restriction are often considered independent barriers to cross-species replication. In this study, we identify a mechanism by which HIV-1 Env adapts to both barriers during SHIV adaptation to macaques: inefficient use of the macaque CD4 receptor and restriction by the type I IFN response. Serial passage of SHIV in macaques selected Env variants with markedly increased resistance to IFN-α, a phenotype that mapped principally to loss of N-linked glycans at N197 and N276 near the CD4-binding site. Removal of these glycans altered the Env conformational landscape, enhanced macaque CD4-dependent entry and membrane fusion, and reduced incorporation of the IFN-inducible restriction factors IFITM3 and IFITM3A into progeny virions. Depletion of endogenous IFITM3/3A eliminated the differential IFN-α sensitivity of parental and adapted viruses, genetically linking passage-associated Env adaptation to escape from IFITM-mediated restriction. Thus, two distinct selective pressures encountered during cross-species adaptation, namely receptor incompatibility and innate immune restriction, can converge on a common set of Env changes that simultaneously improve receptor utilization and promote immune escape.

The acquisition of IFN-α resistance during serial passage is likely to be biologically relevant to SHIV adaptation in vivo. Rapid serial passage is typically performed within the first two weeks of infection, when macaques mount a robust type I IFN response (19–21). Consistent with a role for IFN as a selective pressure during early infection, transmitted/founder HIV-1 variants exhibit increased resistance to type I IFNs, and IFN resistance has been linked to transmission fitness in humans (23, 24). We previously showed that serial passage of SHIVs is accompanied by increasing IFN-α resistance (25), but the viral determinants of this resistance have remained undefined. The strong relationship between replication capacity and IFN-α resistance across Env variants, chimeras, and point mutants suggests that escape from IFN-induced restriction contributes to the increased fitness of adapted SHIVs in macaque cells. Although IFITM escape is unlikely to be the sole determinant of the adapted phenotype, these findings support the idea that the acute type I IFN response imposes selective pressure on Env during adaptation to the macaque host.

A striking feature of this adaptation is that resistance was conferred predominantly by loss of two CD4-binding site-proximal glycans rather than by substitutions at direct CD4-contact residues. N-linked glycans at N197 and N276 are highly conserved, occurring in 97.1% and 95.9% of HIV-1 isolates, respectively (Los Alamos HIV Sequence Database), and contribute to shielding the CD4-binding site, a conserved receptor-binding surface of Env (38, 39, 41). Previous studies have shown that the N197 glycan can restrict accessibility of the CD4-binding site and influence local trimer dynamics, whereas N276 is positioned adjacent to the CD4-binding site (38, 42, 43). Although Env glycans are best known for their role in antibody evasion, glycans at these positions also influence CD4-binding site exposure, Env structure, infectivity, and antigenicity (38, 41, 47–50). In our study, the two glycans made unequal but complementary contributions to adaptation: N197 loss accounted for most of the effects on IFN-α resistance, entry, fusion, and IFITM3/3A incorporation, whereas N276 loss had more modest effects but was required with N197 loss to reproduce the phenotype of the passage-derived Env. Thus, N197 and N276 make non-equivalent but cooperative contributions to Env function. Although loss of these specific glycans may not be selected in all SHIV lineages, our findings illustrate the plasticity of the Env glycan shield as a substrate for functional adaptation.

The preferential enhancement of entry through macaque CD4 provides an important clue to why these changes were selected. Loss of the N197 glycan, alone or together with N276 loss, substantially enhanced macaque CD4-mediated entry without significantly increasing entry through human CD4. Thus, glycan loss does not simply create a globally more infectious Env but appears to compensate for the less favorable interaction between HIV-1 Env and macaque CD4, a major species barrier for most transmitted/founder HIV-1 Envs (10). Previous studies have identified multiple routes to improved macaque CD4 utilization, including substitutions selected during in vitro or in vivo adaptation, engineered changes at residue 375 that enhance macaque CD4 binding, and mutations that favor more open Env conformations (11, 13, 51, 52). These studies show that macaque CD4 usage can be improved not only by substitutions that directly affect receptor engagement but also by changes that alter the conformation of Env. Our findings extend this principle to the glycan shield, showing that the loss of CD4-binding site-proximal glycans can enhance macaque CD4 usage without altering direct CD4-contact residues. Glycan composition therefore represents an additional determinant of species-specific receptor compatibility.

The smFRET results provide a mechanistic bridge between glycan loss and functional adaptation of Env. Removal of the N197 and N276 glycans shifted the conformational ensemble of virus-associated Env, decreasing occupancy of the pre-triggered State 1 and increasing occupancy of the prefusion-closed State 2, with comparatively little change in State 3. These findings indicate that glycan loss does more than increase exposure of the CD4-binding site; it reshapes the conformational landscape sampled by Env. Although we cannot infer that any individual FRET state has increased affinity for macaque CD4, the smFRET data, together with enhanced macaque CD4-dependent entry and membrane fusion, suggest that glycan loss facilitates productive receptor-triggered transitions. One possibility is that the N197 and N276 glycans impose steric or energetic constraints that are readily overcome during engagement of human CD4 but become limiting with the suboptimal macaque receptor; their removal may therefore facilitate downstream conformational transitions required for fusion. Importantly, this differs from a simple model in which macaque adaptation selects a constitutively open Env, because N197/N276 loss did not markedly increase occupancy of the CD4-bound open state. Instead, these glycans appear to tune the Env conformational landscape from which CD4 engagement and membrane fusion proceed.

The strong relationship between membrane fusion and IFN-α resistance further links receptor adaptation to innate immune escape. IFITM proteins are broad-spectrum antiviral restriction factors that inhibit infection by interfering with viral-host membrane fusion, with susceptibility varying among viral glycoproteins (26, 30–33). An Env that engages macaque CD4 more productively and proceeds more efficiently through fusion may therefore be better able to overcome IFITM-mediated restriction, potentially by reducing the opportunity for IFITMs to impede membrane merger. Our results also support a second, non-mutually exclusive mechanism operating in virus-producing cells. Parental SHIV-109mc incorporated more endogenous IFITM3/3A than passage-derived viruses, and transfer of the adapted Env or removal of the N197 glycan was sufficient to reduce virion-associated IFITM3/3A. Because virion-incorporated IFITMs can impair particle infectivity and fusion (28, 29), adaptation may enhance membrane fusion both by improving the intrinsic fusion properties of Env and by reducing IFITM3/3A incorporation into progeny virions. How Env glycan composition controls IFITM incorporation into virions remains unresolved. Glycan-dependent changes in Env could influence its trafficking, localization within producer-cell membranes, partitioning into membrane domains, or organization of viral assembly sites, thereby altering the extent to which IFITMs are incorporated during budding. Although we have no evidence for a direct interaction between Env and IFITM3/3A, the CRISPR depletion experiments establish the functional importance of this association: depletion of endogenous IFITM3/3A rendered parental SHIV-109mc highly IFN-α resistant and abolished differences in IFN-α sensitivity among parental, adapted, and glycan-loss viruses. Thus, IFITM3/3A are major determinants of the Env-dependent IFN phenotype.

These findings also have implications for the interpretation of SHIV models. Macaque adaptation may alter not only CD4 usage but also glycan composition, the Env conformational landscape, membrane fusion, and susceptibility to host restriction factors, properties that should be considered when adapted SHIVs are used to model HIV-1 transmission, neutralization, or vaccine protection. More broadly, our findings extend the functional significance of the HIV-1 glycan shield beyond antibody evasion. Although much attention has focused on antibody-driven evolution of the HIV-1 glycan shield, our findings show that receptor compatibility and innate immune restriction can also shape its evolution. Notably, loss of the N197 and N276 glycans has also been shown to alter susceptibility to neutralizing antibodies (38, 47, 49, 50). This versatility implies a potential evolutionary tradeoff: glycan loss that improves receptor utilization or innate immune escape may also alter exposure of antibody-sensitive Env epitopes. Balancing these competing pressures may influence the maintenance or loss of individual glycans during HIV-1 evolution.

These findings open several avenues for further mechanistic investigation. The smFRET experiments establish that N197/N276 glycan loss can alter conformational sampling of virus-associated Env, but these measurements were performed in a BG505 Env background rather than directly with SHIV-109mc or P3cB Env; the precise redistribution of conformational states may therefore differ among Env backgrounds. The molecular mechanism linking glycan loss to reduced IFITM3/3A incorporation also remains unresolved. In addition, IFITM3 and IFITM3A were analyzed together in the depletion experiments, precluding definitive assignment of their individual contributions. Resolving these effects is further complicated by expansion of the macaque IFITM repertoire and by limited reagents that distinguish closely related macaque IFITM proteins. Future structural, biophysical, and membrane-organization studies should clarify how CD4-binding site-proximal glycans shape Env function and IFITM susceptibility during viral assembly and entry.

Collectively, our findings identify the HIV-1 Env glycan shield as an evolutionary interface at which distinct selective pressures can converge during adaptation to a new host. In macaques, loss of the N197 and N276 glycans simultaneously enhanced utilization of a heterologous CD4 receptor and reduced susceptibility to IFITM3/3A-mediated restriction. Rather than requiring independent solutions to receptor incompatibility and innate immune pressure, cross-species adaptation can therefore select a common set of Env changes that address both barriers. More broadly, this convergence suggests that glycan plasticity may enable lentiviruses to coordinate receptor use, membrane fusion, and resistance to host antiviral pressures during adaptation to a new host.

## Materials and Methods

### Cells, virus stocks, and plasmids

HEK293T cells (ATCC CRL-3216), HEK293T cells stably expressing human or macaque CD4 and CCR5 (53), and HeLa TZM-bl cells (54) (BEI Resources, NIH HIV Reagent Program, catalog no. 8129) were cultured in Dulbecco’s modified Eagle medium (DMEM, Gibco) supplemented with 10% fetal bovine serum (FBS, Sigma), 2 mM L-glutamine (Gibco), and 1x Penicillin-Streptomycin (Gibco) (complete DMEM). Immortalized pig-tailed macaque CD4^+^ T lymphocytes (55) were cultured in Iscove’s modified Dulbecco’s medium (IMDM) supplemented with 10% FBS, 2 mM L-glutamine, and 1x Penicillin-Streptomycin, and 100 U/ml of interleukin-2 (Roche) (complete IMDM).

The SHIV-109mc, SHIV-P3, SHIV-P3N, and SHIV-P4 virus populations have been described previously (35) and were kindly provided by Cecilia Cheng-Mayer.

The full-length proviral SHIV-1157ipd3N4 plasmid has been described previously (8) and was kindly provided by Ruth Ruprecht. Full-length Env expression plasmids derived from the SHIV-P3 and SHIV-P3N virus populations have been described previously (35). Briefly, viral RNA was isolated from plasma, reverse-transcribed, and full-length *env* (gp160) coding sequences were amplified by PCR and cloned into the pCAGGS expression vector using BamHI and EcoRI restriction sites. Six independent Env clones from each virus population were used in this study. Expression plasmids encoding N-terminally HA-tagged macaque IFITM1, IFITM3, IFITM3A, IFITM3-R1, and IFITM3-R2 have been described previously (56). Briefly, the IFITM open reading frames were synthesized as DNA fragments (Twist Bioscience) and cloned into pCMV-HA (Takara Bio) using EcoRI and SalI restriction sites.

### Generation of SHIV infectious molecular clones, Env chimeras, and mutants

The original SHIVC109F.PB4 molecular clone, referred to here as SHIV-109mc, has been described previously (35) and was maintained as separate 5′ SIV and 3′ SHIV hemigenome plasmids. For the present study, a full-length proviral SHIV-109mc infectious molecular clone (IMC) was generated using SHIV-1157ipd3N4 as the proviral backbone. The SHIV-1157ipd3N4 provirus was first transferred into a modified pcDNA3.1(+) backbone using EcoRI and NotI restriction sites. An AfeI-XhoI DNA fragment containing the ZM109F.PB4 *env* open reading frame was synthesized (Genewiz) and inserted into the corresponding AfeI-XhoI region of the SHIV-1157ipd3N4 proviral plasmid by restriction digestion and ligation.

To generate lineage-related SHIV IMCs, three HIV-1 Env variants from each of the P3 and P3N virus populations were introduced into the SHIV-1157ipd3N4 proviral backbone using a two-step cloning strategy. Briefly, the AfeI-XhoI fragment of SHIV-1157ipd3N4 encompassing the *env*-containing region was cloned into a modified pcDNA3.1(+) backbone to generate an intermediate plasmid. P3 and P3N *env* variants were then introduced into the intermediate plasmid using KpnI and XhoI restriction sites, after which the corresponding AfeI-XhoI fragment was transferred back into the SHIV-1157ipd3N4 proviral plasmid to generate replication-competent P3- and P3N-derived SHIV IMCs. This strategy generated P3- and P3N-derived SHIV IMCs that differed only in their HIV-1 Env sequences while retaining an otherwise identical proviral backbone. The resulting panel included three independent P3 Env variants (P3cA, P3cB, and P3cC) and three independent P3N Env variants (P3NcA, P3NcB, and P3NcC).

Chimeras between SHIV-109mc and SHIV-P3cB were generated by replacing the AfeI-XhoI region of the SHIV-109mc proviral plasmid with synthesized DNA fragments (Twist Bioscience) encoding the indicated SHIV-P3cB Env gp120, gp41, or gp120 subregions. SHIV-109mc constructs encoding the indicated amino acid substitutions within the C2 region were generated using the same DNA fragment synthesis and restriction enzyme-based cloning strategy.

The full-length SHIV-109mc and the P3- and P3N-derived SHIV proviral plasmids were verified by complete plasmid sequencing, and Env chimeras and point-mutant constructs were verified by DNA sequencing of the entire *env* gene.

### SHIV production

Stocks of SHIV-109mc, SHIV-P3, SHIV-P3N, and SHIV-P4 virus populations were expanded for 6 days in immortalized macaque CD4^+^ T lymphocytes, as described previously (25). Replication-competent SHIVs were generated by transfecting 2×10^6^ HEK293T cells with 6 μg of SHIV IMC plasmid DNA using Fugene 6 transfection reagent (Roche) following manufacturer’s protocol. At 48 hours post-transfection, virus-containing supernatants were harvested, passed through a 0.2-μm sterile filter, and concentrated approximately 10-fold using Amicon Ultracel 100 kDa centrifugal filters (Millipore). SHIV titers were determined by infection of TZM-bl cells followed by quantification of β-galactosidase activity at 48 hours post-infection, as described previously (54).

### SHIV replication assays

SHIV replication kinetics were assessed as described previously (34). Briefly, 4×10^6^ immortalized macaque CD4^+^ T lymphocytes were infected at an MOI of 0.02 by spinoculation at 1200 x g for 90 minutes at room temperature. Following spinoculation, cells were washed four times with 1 mL complete IMDM, resuspended in 5 mL complete IMDM, and cultured in a single well of a 6-well plate. Every 3 days, two-thirds of each culture was removed and replaced with fresh complete IMDM. Cell-free viral supernatants were collected from the harvested cultures by centrifugation at 650 x g for 5 minutes at room temperature. Viral replication was quantified by measuring reverse transcriptase (RT) activity in culture supernatants using the RT activity assay.

### IFN-α sensitivity assays

SHIV sensitivity to IFN-α was assessed as described previously (25). Briefly, 4.25×10^6^ immortalized macaque CD4^+^ T lymphocytes were infected with each SHIV at an MOI of 0.02 by spinoculation at 1200 x g for 90 minutes at room temperature. Following spinoculation, cells were washed three times with 1 mL complete IMDM and plated at 2.5×10^5^ cells per well in black 96-well polystyrene plates containing 250 μL complete IMDM supplemented with 0–5,000 U/mL IFN-α2a (PBL Assay Science) in duplicate. At 3 days post-infection, culture supernatants were removed and replaced with fresh complete IMDM containing the corresponding concentration of IFN-α2a. Supernatants were harvested at 6 days post-infection, and infectious virus production was quantified using TZM-bl indicator cells and the Galacto-Light Plus β-Galactosidase Assay (ThermoFisher) following manufacturer’s protocol. Percent viral inhibition was calculated relative to matched untreated controls as 100 x [1 − (β-galactosidase activity in the IFN-α2a-treated sample/β-galactosidase activity in the untreated sample)]. IFN-α IC_50_ values, defined as the concentration of IFN-α2a resulting in 50% inhibition of viral replication, were determined from a nonlinear, best-fit curve using GraphPad Prism software.

### Phylogenetic and sequence analyses

Env nucleotide sequences were aligned and analyzed using Geneious Prime software. Phylogenetic relationships were inferred using the neighbor-joining method with the Tamura-Nei genetic distance model, and branch support was assessed using 1,000 bootstrap replicates. Env amino acid sequences were aligned using the MUSCLE algorithm in Geneious Prime.

### Structural modeling

A homology model of SHIV-109mc Env was generated using the BG505 Env trimer (PDB 5FYL) and ZM109F.PB4 gp120 core (PDB 3TIH) structures as templates using HHPred software tools (57). The SHIV-109mc Env model was structurally superimposed with the BG505 Env atomic model to compare the positions corresponding to N197 and N276. Structures of gp120 bound to human CD4 (PDB 1GC1) and macaque CD4 (PDB 6TYB) were then superimposed with the Env model to examine the spatial relationship of the N197- and N276-linked glycans, which are resolved in the BG505 atomic structure, to the CD4-binding interface. All structural comparisons and superimpositions were performed in UCSF Chimera using the MatchMaker tool (58).

### Preparation and fluorescent labeling of virus-associated Env for smFRET

Replication-incompetent, reverse transcriptase-deficient (ΔRT) HIV-1_Q23_ virions bearing BG505 Env with the N197D and N276D substitutions were generated and fluorescently labeled for smFRET imaging as described previously (44), with modifications. A dual-amber BG505 Env construct bearing the N197D and N276D substitutions and containing TAG codons at positions N136 in V1 and S401 in V4 (N136^TAG^/S401^TAG^) was expressed using an amber-suppression system to enable site-specific incorporation of noncanonical amino acids (ncAAs). The resulting N136^ncAA^/S401^ncAA^ Env was subsequently labeled with fluorophores by bioorthogonal click chemistry.

Viral particles were produced by cotransfecting HEK293T cells with an amber-free HIV-1_Q23_ Env_BG505_ ΔRT plasmid, an HIV-1_Q23_ Env_BG505_ N197D/N276D N136^TAG^/S401^TAG^ ΔRT plasmid, and the amber-suppression plasmid tRNAPyl/NES-PylRS^AF^ (59). The amber-suppression system provided the cognate tRNA and aminoacyl-tRNA synthetase required for site-specific incorporation of the ncAA at the engineered TAG codons. Transfections were performed in the presence of 250 μM TCO*A (trans-cyclooct-2-en-L-lysine; SiChem, catalog no. SC8008), a click-reactive ncAA. The relative amounts of amber-free and dual-TAG Env plasmids were optimized based on previously characterized Env expression and incorporation efficiencies (44) to favor incorporation of, on average, a single dual-ncAA-containing Env protomer per Env trimer on individual viral particles.

At 40 hours post-transfection, viral particles were concentrated by ultracentrifugation at 25,000 rpm for 2 hours at 4°C. Viral pellets were resuspended in labeling buffer containing 50 mM HEPES, 10 mM MgCl_2_, and 10 mM CaCl_2_. Site-specific fluorescent labeling was performed as described previously (44). TCO*A residues incorporated at N136 and S401 were conjugated to the tetrazine-functionalized fluorophores LD555-TTZ and LD655-TTZ, respectively, at a final concentration of 0.1 μM by bioorthogonal click chemistry. The fluorophores were custom synthesized by Lumidyne Technologies. Labeling reactions were carried out for approximately 6 hours at room temperature in the dark.

For subsequent surface immobilization during smFRET imaging, PEG2000-biotin was added to a final concentration of 0.1 mg/mL, and labeled viral particles were incubated for 30 minutes at room temperature with gentle rotation. Labeled virions were then purified by density-gradient ultracentrifugation through a 6–18% OptiPrep gradient at 40,000 rpm for 1 hour at 4°C.

### smFRET data acquisition and analysis

smFRET measurements of fluorescently labeled virus-associated Env were performed as described previously (44, 46, 60). Briefly, fluorescently labeled HIV-1_Q23_ virions bearing BG505 Env with the N197D and N276D substitutions were immobilized in a PEG-passivated, biotinylated quartz imaging chamber functionalized with streptavidin. A 532-nm laser was coupled through a prism to generate an evanescent excitation field at the quartz–aqueous interface. Donor fluorophores were excited by the evanescent field, and donor and acceptor fluorescence emissions were spectrally separated and recorded simultaneously at 25 Hz for 80 seconds (2,000 frames).

Image sequences were analyzed using SPARTAN software (61) and custom MATLAB scripts. Individual donor and acceptor fluorescence trajectories were extracted from the image stacks, followed by background subtraction and correction for donor-to-acceptor spectral cross-talk. FRET efficiency was calculated for each frame as FRET = I_A_/(γI_D_ + I_A_), where I_D_ and I_A_ denote the background-corrected donor and acceptor fluorescence intensities, respectively, and γ is a correction factor accounting for differences in the relative detection efficiencies of the two fluorescence channels.

Individual FRET trajectories were subjected to predefined quality-control criteria and retained if they exhibited a single photobleaching event, an adequate signal-to-noise ratio, and anticorrelated donor and acceptor intensity changes. These criteria were used to exclude particles containing multiple fluorescently labeled Env protomers or multiple labeled Env molecules, particles lacking detectable labeling, and trajectories arising from inactive or incompletely labeled Env molecules. Automated filtering was complemented by manual inspection of individual trajectories.

Selected FRET trajectories were idealized using a three-state hidden Markov model (HMM) (62). Trajectories satisfying all selection criteria were pooled to generate FRET efficiency histograms. FRET distributions were fitted in MATLAB by least-squares fitting to the sum of three Gaussian components corresponding to distinct Env conformational states. The relative occupancy of each conformational state was calculated from the integrated area of the corresponding Gaussian component and expressed as the mean probability of Env occupying that conformational state.

### HIV-1 Env-mediated entry assays

Green fluorescent protein (GFP)-reporter HIV-1 pseudoviruses were generated as described previously (52). Briefly, HEK293T cells were cotransfected with 4 µg of Envelope (Env)-deficient HIV-1 proviral plasmid (Q23ΔEnvGFP) and 2 µg of the indicated Env expression plasmid using Fugene 6 transfection reagent (Roche) following manufacturer’s protocol.

Entry assays in HEK293T cells expressing human or macaque CD4 and CCR5 were performed as described previously (52). Briefly, cells were infected with GFP-reporter pseudoviruses at an MOI of 1 in the presence of 10 μg/mL DEAE-dextran by spinoculation at 1200 x g for 90 minutes. At 48 hours post-infection, cells were harvested, fixed in 2% paraformaldehyde, washed twice, and resuspended in fluorescence-activated cell sorter (FACS) buffer (1x phosphate-buffered saline (PBS), 1% FBS, 1 mM EDTA). GFP expression was measured using Attune NxT flow cytometer, and data were analyzed using FlowJo software.

### SHIV membrane fusion assays

The SHIV β-lactamase-Vpr (BlaM-Vpr) fusion assay was performed as described previously (52). BlaM-SIVmac239 Vpr-containing SHIV particles were generated by co-transfecting HEK293T cells with 4.5 μg of proviral SHIV plasmid and 1.5 μg of pcDNA3.1-BlaM-SIVmac239 Vpr plasmid using Fugene 6 transfection reagent following manufacturer’s protocol. At 48 hours post-transfection, virus-containing supernatants were harvested, passed through a 0.2-μm sterile filter, concentrated approximately 10-fold using Amicon Ultracel 100 kDa centrifugal filters (Millipore), aliquoted, and stored at −80°C. RT activity of viral stocks was measured using the RT activity assay.

For fusion assays, 1×10^5^ immortalized macaque CD4^+^ T lymphocytes in 100 μL complete IMDM were infected with BlaM-SIVmac239 Vpr-containing SHIVs equivalent to 500 mU of RT by spinoculation at 1200 x g for 90 minutes at room temperature, followed by incubation for 1 hour at 37°C in 5% CO_2_. SHIV fusion was quantified by monitoring cleavage of the fluorescent CCF2-AM substrate. Following infection, cells were washed once with 300 μL cold CO_2_-independent medium (Gibco) without FBS and resuspended in 100 μL CO_2_-independent medium supplemented with 10% FBS. Cells were then loaded with CCF2-AM substrate (LiveBLAzer CCF2-AM Kit, Invitrogen) following manufacturer’s protocol in the presence of 1.8 mM probenecid (Sigma) and incubated for 2 hours. Cells were washed three times with 300 μL cold CO_2_-independent medium without FBS and once with 1x PBS, fixed in 200 μL of 2% paraformaldehyde, washed once with 1x PBS, and resuspended in 400 μL FACS buffer. Fluorescence corresponding to cleaved CCF2-AM substrate was measured on Attune NxT flow cytometer, and data were analyzed using FlowJo software.

### CRISPR-Cas9-mediated depletion of IFITM3/3A

IFITM knockout cell pools were generated by electroporation of CRISPR-Cas9 ribonucleoprotein (crRNP) complexes, as described previously (34), with modifications. Immortalized macaque CD4^+^ T lymphocytes were electroporated with custom CRISPR RNAs (crRNAs) targeting IFITM3 and IFITM3A or with a non-targeting control (NTC) crRNA. The crRNA sequences were as follows: IFITM3, 5′-GAGCACGAGGUGGCUAUGAU-3′; IFITM3A, 5′-GCUGGCCGCUGUUGACAGGA-3′; and NTC, 5′-UAGACAACCGCGGAGAAUGC-3′.

crRNAs (Integrated DNA Technologies) and tracrRNA (Integrated DNA Technologies, catalog no. 1072534) were resuspended at 160 μM in 10 mM Tris-HCl pH 7.4. For crRNP assembly, 1 μL crRNA was mixed at an equimolar ratio with 1 μL tracrRNA and incubated for 30 minutes at 37°C. Cas9-NLS protein (2 μL of a 40 μM stock, UC Berkeley MacroLab) was then added, and the mixture was incubated for an additional 15 minutes at 37°C to generate IFITM3/3A-targeting or NTC crRNP complexes.

For each electroporation, 3.5 μL crRNP complex was added to 5×10^5^ macaque CD4^+^ T lymphocytes resuspended in SG Cell Line Nucleofector Solution (Lonza, catalog no. V4SC-3096), and cells were electroporated using a Lonza 4D-Nucleofector according to the manufacturer’s protocol. Immediately after electroporation, 80 μL prewarmed complete IMDM was added, and cells were allowed to recover for 30 minutes at 37°C. Forty-eight replicate electroporations were performed for each crRNA condition. Following recovery, cells from replicate electroporations were pooled, resuspended at 1.6×10^6^ cells/mL in complete IMDM, and cultured in 6-well plates. Depletion of IFITM3/3A was assessed by immunoblotting at 7 days post-electroporation.

### Immunoblotting and virion incorporation assays

Whole cell extracts were prepared by lysing the cells in radioimmunoprecipitation assay (RIPA) cell lysis buffer (50 mM Tris pH 8.0, 0.1% SDS, 1% Triton-X, 150 mM NaCl, 1% deoxycholic acid, 2 mM PMSF). For analysis of IFITM incorporation into virions, virus-containing supernatants from the infected or transfected cell cultures were centrifuged at 650 x g for five minutes at room temperature. Cell-free supernatant was filtered through a 0.2 µm filter and then pelleted through a 25% sucrose cushion by ultracentrifugation for at 28,000 rpm for 90 minutes at 4°C. Virus pellets were resuspended in 70 μL RIPA buffer and incubated for 10 minutes at room temperature. Virion lysates were normalized based on RT activity, and equivalent amounts of lysate were subjected to SDS-PAGE and immunoblotted. Standard Western blotting procedures were used with the following antibodies: IFITM3 (Proteintech, catalog no. 11714-1-AP), SIV p27 (ABL, catalog no. 4323), HA.11 epitope tag (BioLegend, catalog no. 901521), and β-Actin (Abcam, catalog no. ab3280). The IFITM3 antibody used in this study recognizes both macaque IFITM3 and IFITM3A, as described previously (34); accordingly, immunoblot signals detected with this antibody are designated as “IFITM3/3A.”

### Reverse transcriptase activity assays

Reverse transcriptase (RT) activity assay was performed as described previously (63). Briefly, 5 μL of viral supernatant, virus stock, or RT standard was mixed with 5 μL of 2x lysis buffer (100 mM Tris-HCl, pH 7.4, 50 mM KCl, 0.25% Triton X-100, and 40% glycerol) containing 4 U RNaseOUT (Invitrogen) and incubated for 10 minutes at room temperature. Viral lysates were then diluted 1:10 with 90 μL nuclease-free water (Life Technologies). For each qRT-PCR reaction, 9.6 μL of diluted viral lysate was combined with 10.4 μL of reaction mixture containing 10 µl of 2x Maxima SYBR Green/ROX qPCR Master Mix (ThermoFisher), 0.1 µl of 4U/µl RNaseOUT, 0.1 µl of 0.8 µg/µl MS2 RNA template (Roche), and 0.1 µl each of 100 µM forward 5’-TCCTGCTCAACTTCCTGTCGAG-3’ and reverse 5’-CACAGGTCAAACCTCCTAGGAATG-3’ primers. qRT-PCR was performed using a QuantStudio 3 Real-Time PCR System (Applied Biosystems). RT activity was quantified from a standard curve generated using recombinant reverse transcriptase (Millipore, catalog no. 382129).

## Figure Legends

**Figure S1. Env sequences recovered after SHIV passage in macaques form distinct lineages.** Neighbor-joining phylogeny of full-length Env nucleotide sequences from SHIV-109mc and Env clones recovered from the P3 and P3N virus populations. SHIV-109mc Env is shown in blue, P3 Env clones in orange, and P3N Env clones in red. Bootstrap support values ≥50 are shown at internal nodes. The scale bar indicates 0.002 substitutions per site.

**Figure S2. Env sequences recovered after SHIV passage in macaques acquire multiple amino acid substitutions.** Amino acid alignment of SHIV-109mc Env and Env sequences recovered from the P3 and P3N virus populations. Dots indicate amino acid identity with SHIV-109mc Env, and substitutions are shown using single-letter amino acid notation. Env residue numbers are indicated above the alignment.

**Figure S3. SHIV replication kinetics correlate with IFN-**α **resistance. (A)** Replication kinetics of SHIV-109mc and infectious molecular clones encoding the indicated P3 or P3N Envs in immortalized macaque CD4^+^ T lymphocytes. Supernatant reverse transcriptase activity was measured at the indicated days postinfection. Data are means from three independent experiments; error bars indicate standard deviations. **(B)** Relationship between replication kinetics and IFN-α resistance. The slope of each replication curve through day 6 was plotted against the corresponding IFN-α IC_50_ value. The association was evaluated by Spearman rank correlation (*r*_s_ = 0.9870, *P* < 0.0001).

**Figure S4. SHIV replication kinetics correlate with IFN-**α **resistance across Env chimeras and glycan mutants. (A, C, and E)** Replication kinetics of the indicated parental SHIVs, Env chimeras, and SHIV-109mc point mutants in immortalized macaque CD4^+^ T lymphocytes. Supernatant reverse transcriptase activity was measured at the indicated days postinfection. Data are means from three independent experiments; error bars indicate standard deviations. **(B, D, and F)** The slope of each replication curve through day 6 was plotted against the corresponding IFN-α IC_50_ value. Associations were evaluated by Spearman rank correlation for the gp120/gp41/C2-V4 chimera panel (B; *r*_s_ = 0.9795, *P* = 0.0013), the individual gp120 subregion chimera panel (D; *r*_s_ = 0.6154, *P* = 0.0647), and the SHIV-109mc point-mutant panel (F; *r*_s_ = 0.9660, *P* < 0.0001).

**Figure S5. Loss of N197 and N276 glycans does not alter human CD4-dependent entry.** Entry mediated by the indicated Envs in HEK293T cells expressing human CD4 and CCR5. GFP-reporter viruses pseudotyped with each Env were used to infect target cells, and entry was quantified as the percentage of GFP-positive cells by flow cytometry. Data are means from four independent experiments; error bars indicate standard deviations. Statistical significance was determined by ordinary one-way ANOVA followed by Dunnett’s multiple-comparisons test relative to 109mc Env; no significant differences were detected.

**Figure S6. Representative flow cytometry analysis of SHIV membrane fusion.** Representative flow-cytometry plots from the BlaM-Vpr fusion assay quantified in Fig. 3H. Uninfected cells and cells infected with SHIV-109mc, SHIV-P3cB, the C2 chimera, or the indicated SHIV-109mc point mutants are shown. Cells containing cleaved CCF2-AM substrate are identified within the indicated gate, and the percentage of fusion-positive cells is shown for each condition.

**Figure S7. Virion incorporation of IFITM3 and IFITM3A differs between parental and passaged SHIVs. (A)** Expression of HA-tagged macaque IFITM proteins in HEK293T producer cells. Cells were transfected with empty vector or plasmids encoding the indicated HA-tagged macaque IFITM proteins, and lysates were analyzed by immunoblotting for HA and β-actin. **(B)** Incorporation of macaque IFITM proteins into SHIV-109mc and SHIV-P3cB particles. Virions were produced in HEK293T cells coexpressing the indicated HA-tagged IFITM proteins, normalized by reverse transcriptase activity, and analyzed by immunoblotting for HA and SIV p27.

**Table S1. Parameters for three-state modeling of FRET histograms and estimation of conformational state occupancies.** FRET histograms for virus-associated HIV-1 BG505 Env bearing the N197D and N276D substitutions and the corresponding wild-type BG505 Env reference from our previous study (44) were independently fitted using constrained three-state models. Each conformational state, State 1 (pre-triggered, PT; centered at ∼0.1 FRET), State 2 (prefusion closed, PC; centered at ∼0.3 FRET), and State 3 (CD4-bound open, CO; centered at ∼0.6 FRET), was modeled as a Gaussian distribution, N(μ, σ²). State occupancies are reported as mean probabilities. Model fit was assessed using the coefficient of determination (R²) and root mean square error (RMSE), with values approaching 1 and 0, respectively, indicating better fits.

## Acknowledgments

We thank Cecilia Cheng-Mayer for providing SHIV-109mc, SHIV-P3, SHIV-P3N, and SHIV-P4 virus populations and P3- and P3N-derived Env clones; Ruth Ruprecht for providing the SHIV- 1157ipd3N4 plasmid; and Hans-Peter Kiem for providing immortalized macaque CD4^⁺^ T lymphocytes. TZM-bl cells were obtained through BEI Resources, NIAID, NIH.

## Funding

This work was supported by grants from the National Institutes of Health (NIAID R00 AI125136, R56 AI172615, R01 AI172615, and a U54 AI170752 Collaborative Development Award to AS, NIDA DP1 DA039543 and NICHD R01 HD103571 to JO, and NIAID R01 AI181600 to ML).

## Competing interests

The authors declare no competing interests.

## Data Availability

The data supporting the findings of this study are provided in the main text, figures, and Supplementary Information. Additional data are available from the corresponding author upon reasonable request. Previously published structures used in this study include PDB IDs: 5FYL, 3TIH, 1GC1, 6TYB, 4ZMJ, and EMD ID: 21412.

## Author contributions

A.S. and J.O. conceived and designed the study. A.S. supervised and oversaw the research. A.C.S., H.W., and D.K. performed the virological and cell-based experiments. V.M.P. conducted structural modeling. R.H.K. and M.L. performed and analyzed the smFRET experiments. A.C.S., A.P., and A.S. analyzed and interpreted the data. A.S. wrote the original manuscript. All authors reviewed and edited the manuscript and approved the final version.

## Notes

### Competing Interest Statement

The authors have declared no competing interest.

