## Supplemental Figures S1-S7 for "HIV-1 envelope glycan loss links receptor adaptation and innate immune escape in SHIVs"

### Slide 1
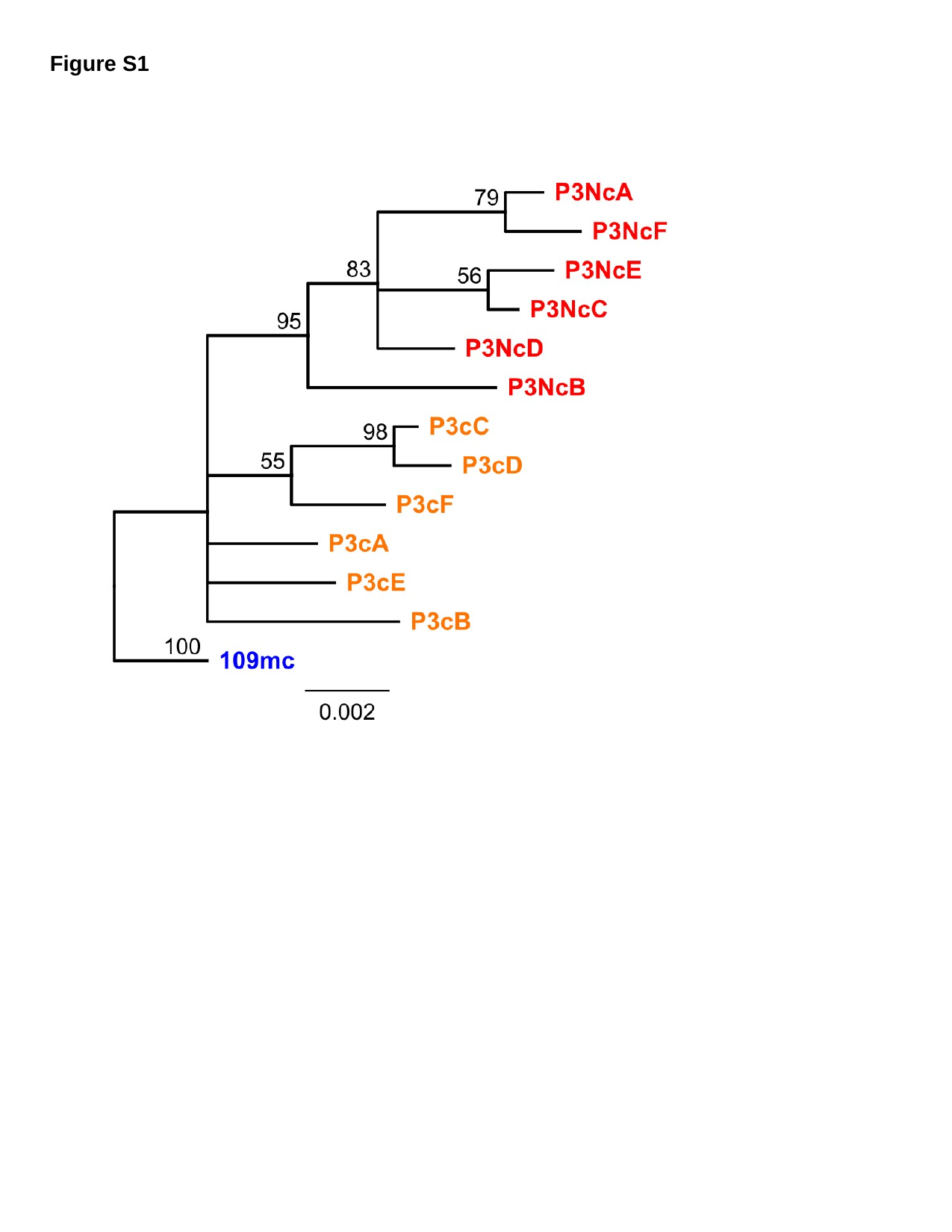

Figure S1

### Slide 2
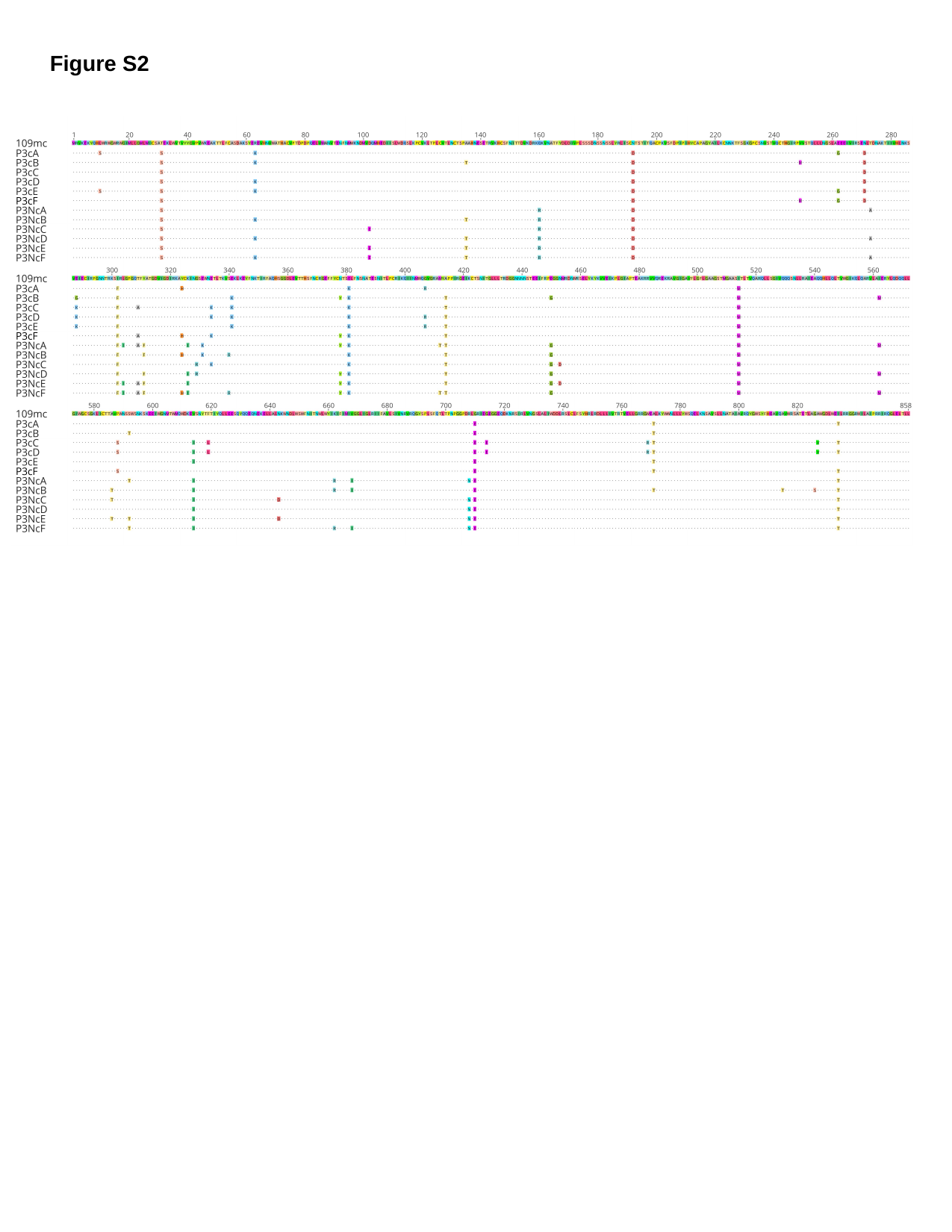

Figure S2

### Slide 3
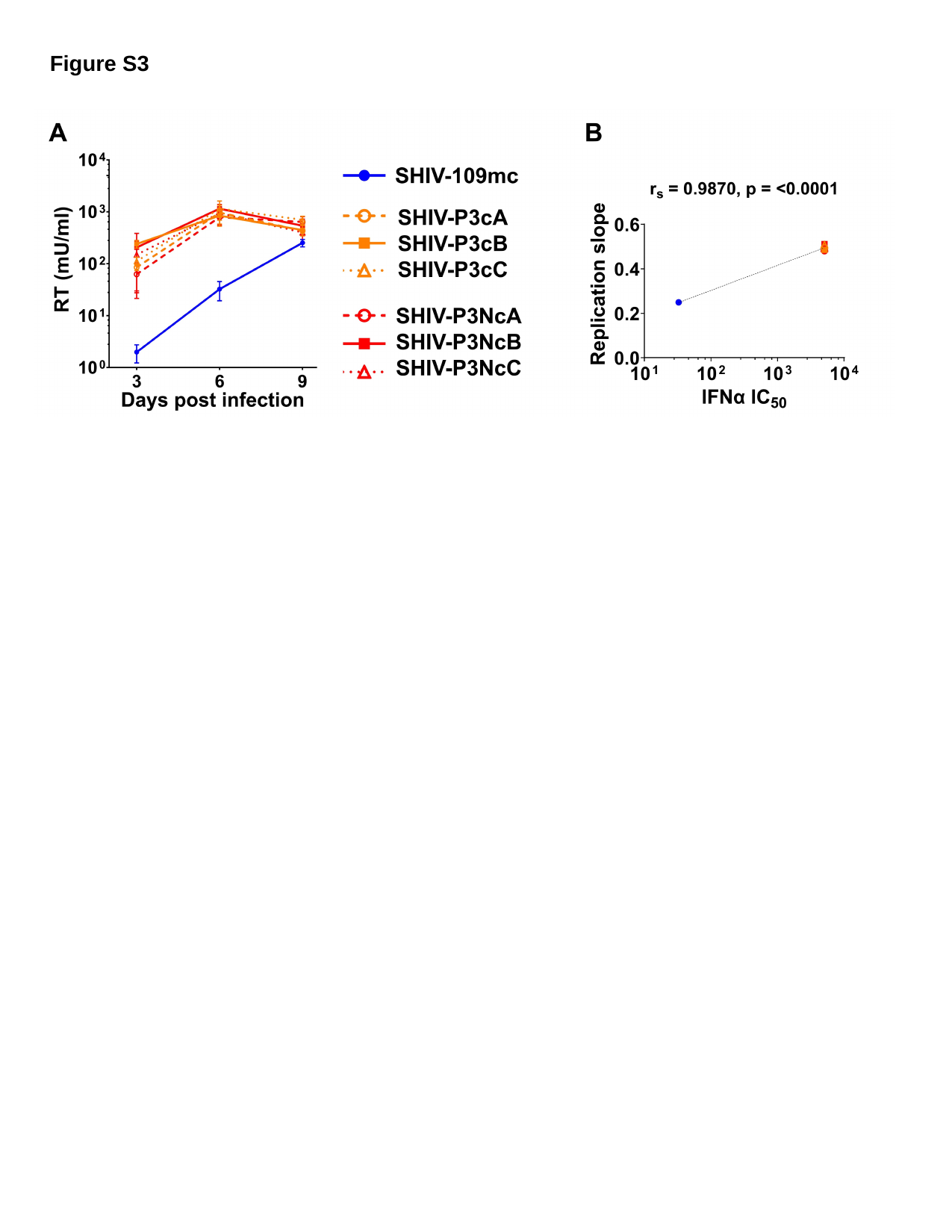

Figure S3

### Slide 4
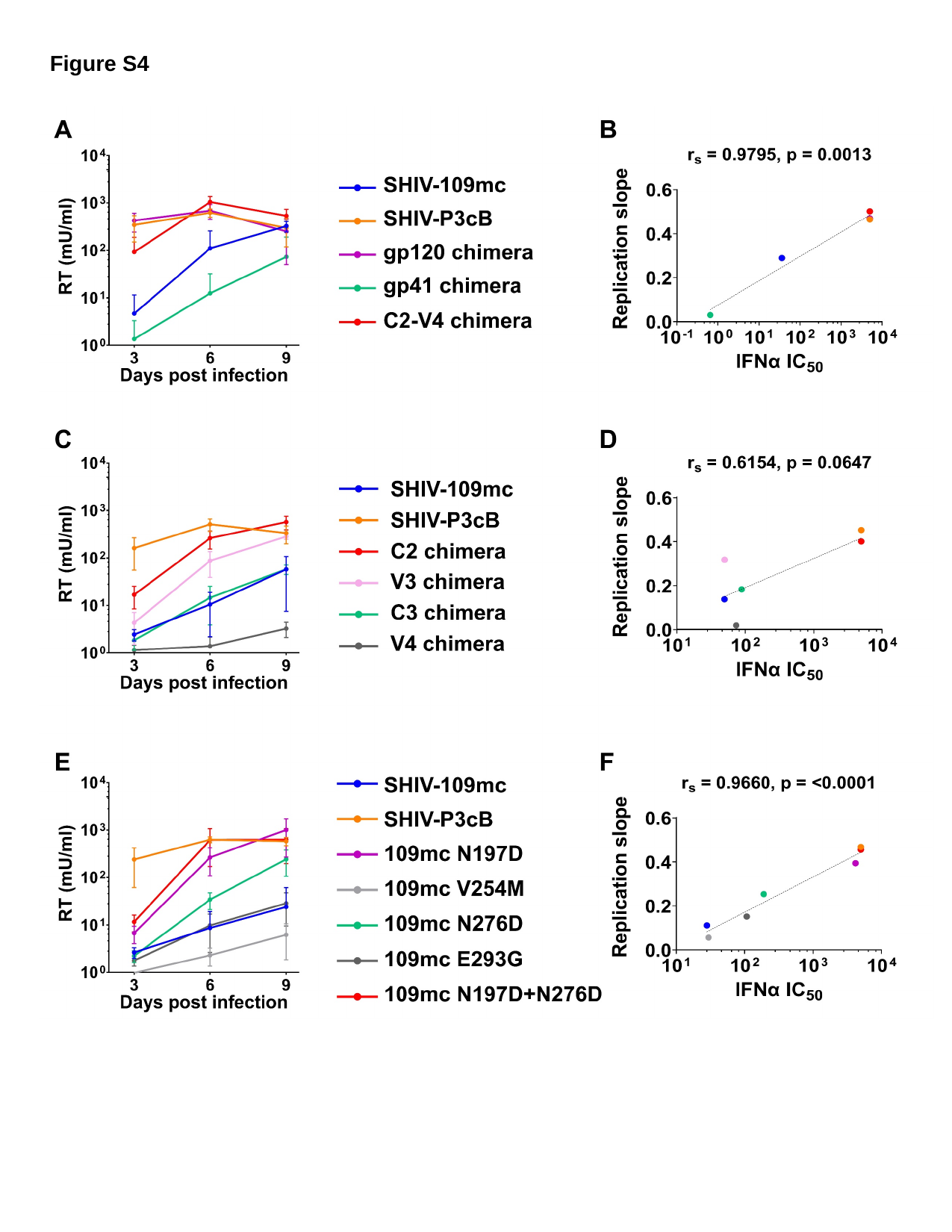

Figure S4

### Slide 5
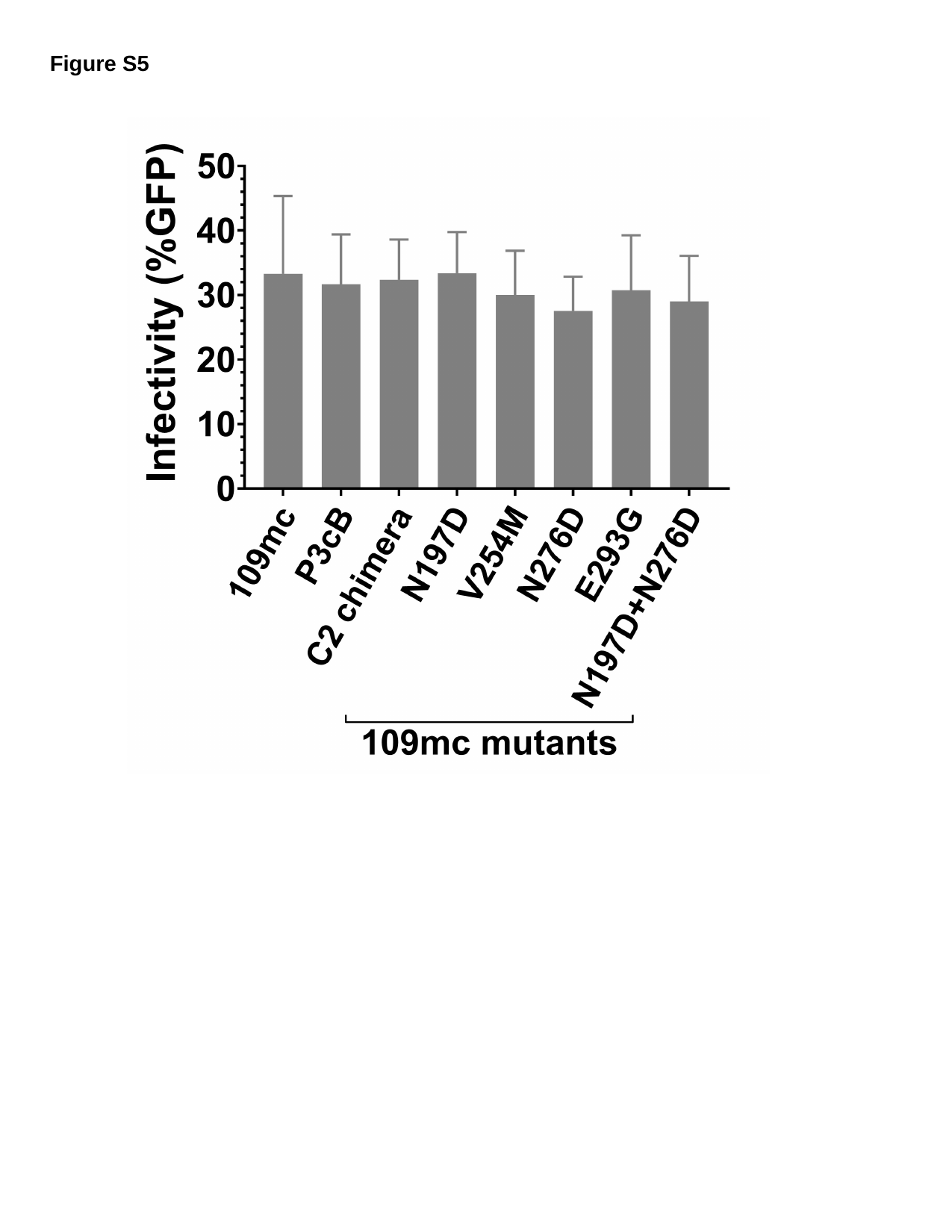

Figure S5

### Slide 6
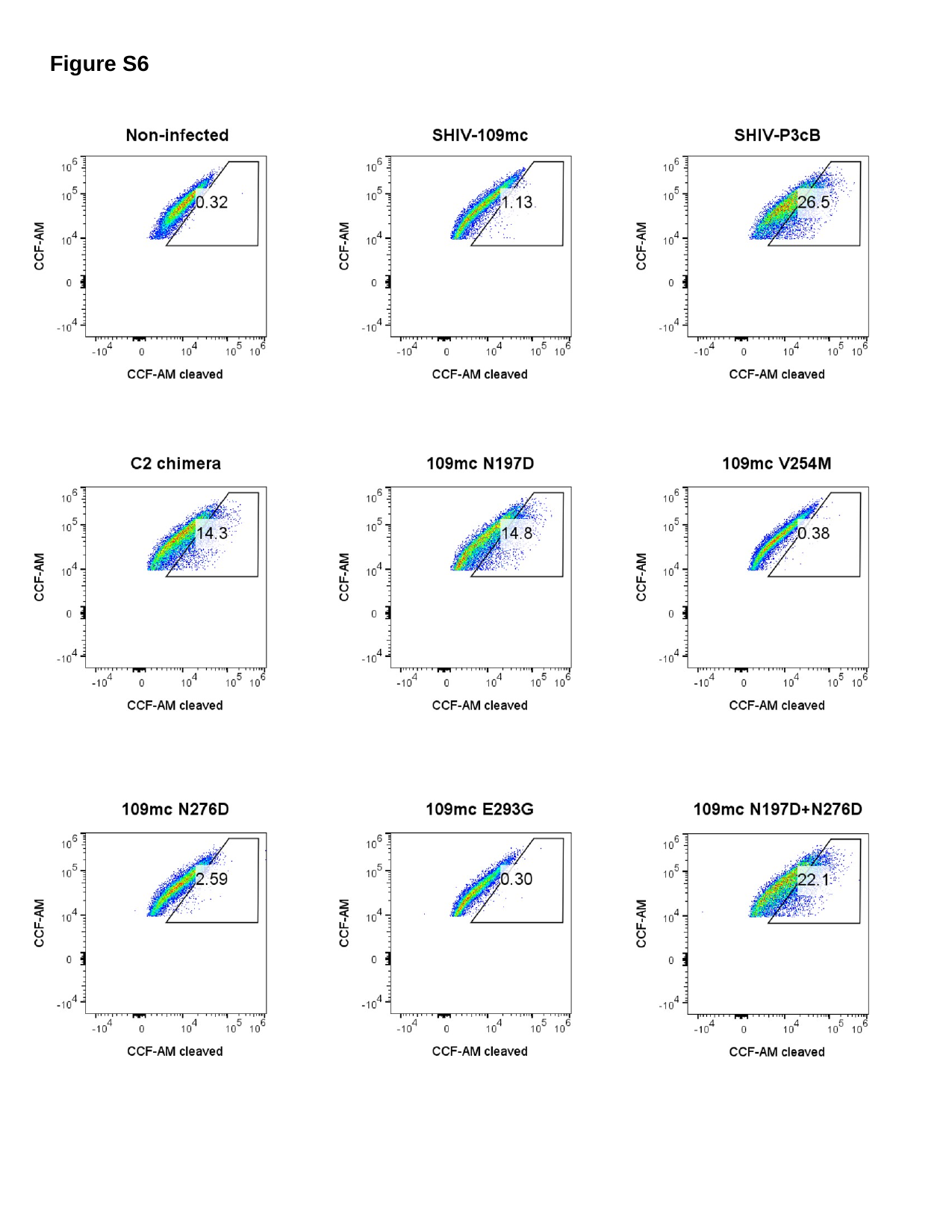

Figure S6

### Slide 7
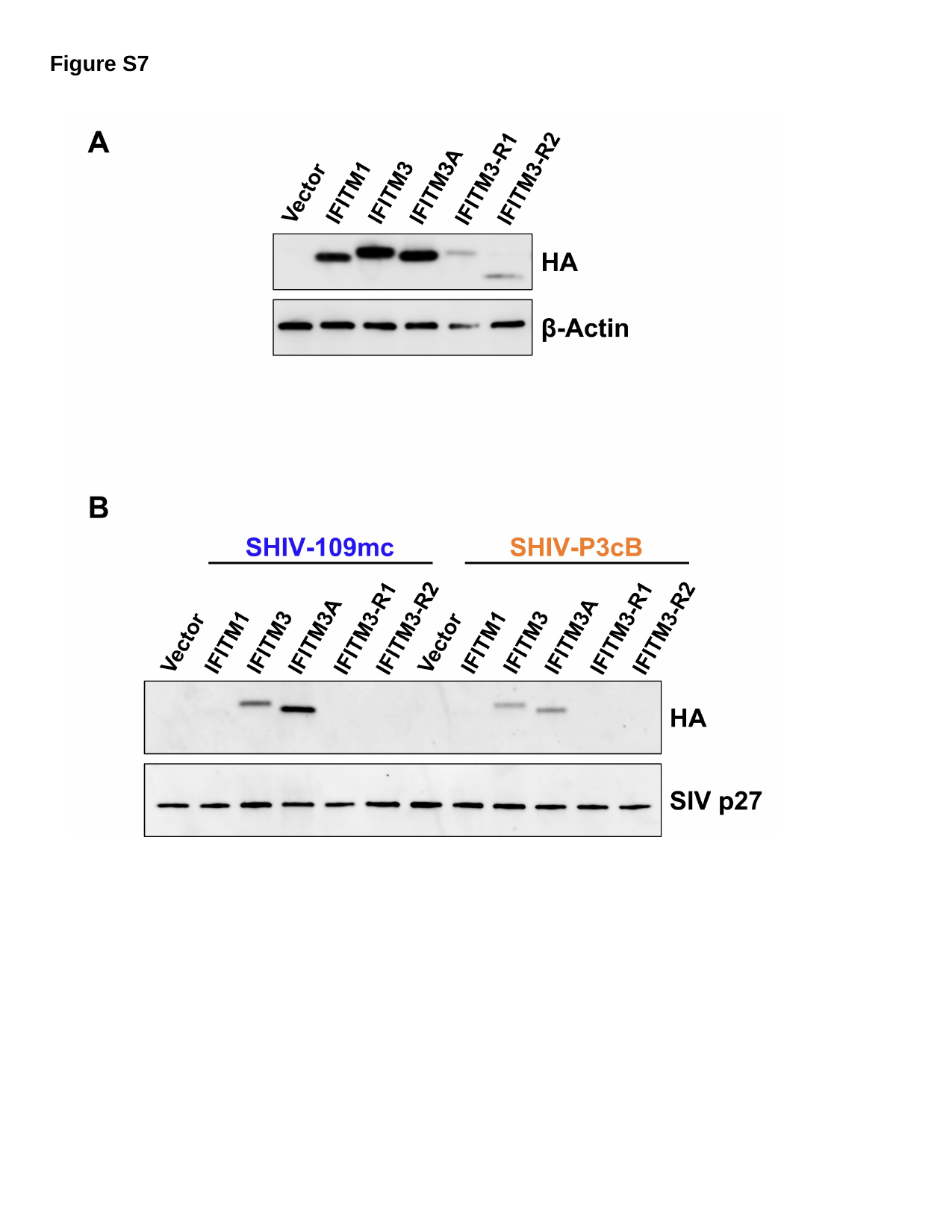

Figure S7
